# Bayesian Inference of Binding Kinetics from Fluorescence Time Series

**DOI:** 10.1101/2025.02.03.636267

**Authors:** J. Shepard Bryan, Stanimir Asenov Tashev, Mohamadreza Fazel, Michael Scheckenbach, Philip Tinnefeld, Dirk-Peter Herten, Steve Pressé

## Abstract

The study of binding kinetics via the analysis of fluorescence time traces is often confounded by measurement noise and photophysics. Although photoblinking can be mitigated by using labels less likely to photoswitch, photobleaching generally cannot be eliminated. Current methods for measuring binding and unbinding rates are therefore limited by concurrent photobleaching events. Here, we propose a method to infer binding and unbinding rates alongside photobleaching rates using fluorescence intensity traces. Our approach is a two-stage process involving analyzing individual regions of interest (ROIs) with a Hidden Markov Model to infer the fluorescence intensity levels of each trace. We then use the inferred intensity level state trajectory from all ROIs to infer kinetic rates. Our method has several advantages, including the ability to analyze noisy traces, account for the presence of photobleaching events, and provide uncertainties associated with the inferred binding kinetics. We demonstrate the effectiveness and reliability of our method through simulations and data from DNA origami binding experiments.

## 1 Introduction

The study of binding kinetics is critical to understanding the behavior of biological systems and chemical reactions at the molecular level. For instance, accurate determination of binding kinetics can provide valuable insights into the mechanisms of protein self-assembly [1, 2, 3, 4], enzymatic and protein-protein interactions [5, 6, 7, 8, 9, 10], DNA-protein interactions and gene expression [11, 12, 13], nuclear breakdown during mitosis [14], nuclear pore complex (NPC) assembly [15, 16, 17], and the signaling mechanism and cell response via receptor clustering and protein recruitment [18, 19] among others. Likewise, binding rates are essential to understand and control reactions in chemistry, including but not limited to nanocatalysis [20, 21], polymerization catalysis [22] and fundamental reaction mechanisms [23, 24, 25]. Currently, several techniques exist for measuring binding kinetics, including surface plasmon resonance [26, 27], and fluorescence correlation spectroscopy [28, 29, 30, 31], among others [32]. However, these methods have limitations, such as the difficulty in analyzing data in the low signal-to-noise ratio (SNR) regime or their inability to account for photophysical events including photobleaching, or simply termed bleaching, which may be confounded with unbinding.

Quantifying the stoichiometric fluctuations of clusters [33, 34, 35, 36, 37, 38, 39] is another important motivation for studying binding kinetics. Here, super-resolution methods such as PALM, dSTORM, and qPAINT provide means to estimate the stoichiometry of proteins clusters [40, 41, 42, 43] albeit mostly limited to static structures, *e*.*g*., fixed cells. Hidden Markov models [44] can be used to infer transition probabilities, but in the presence of photophysics, these do not directly translate to rates which becomes relevant when studying faster events. Moreover, current methods for inferring binding kinetics are prone to deviations, such as overestimating unbinding probabilities, in their estimates due to binding kinetics and photophysical effects, such as photobleaching (figure 1).

**Figure 1:**
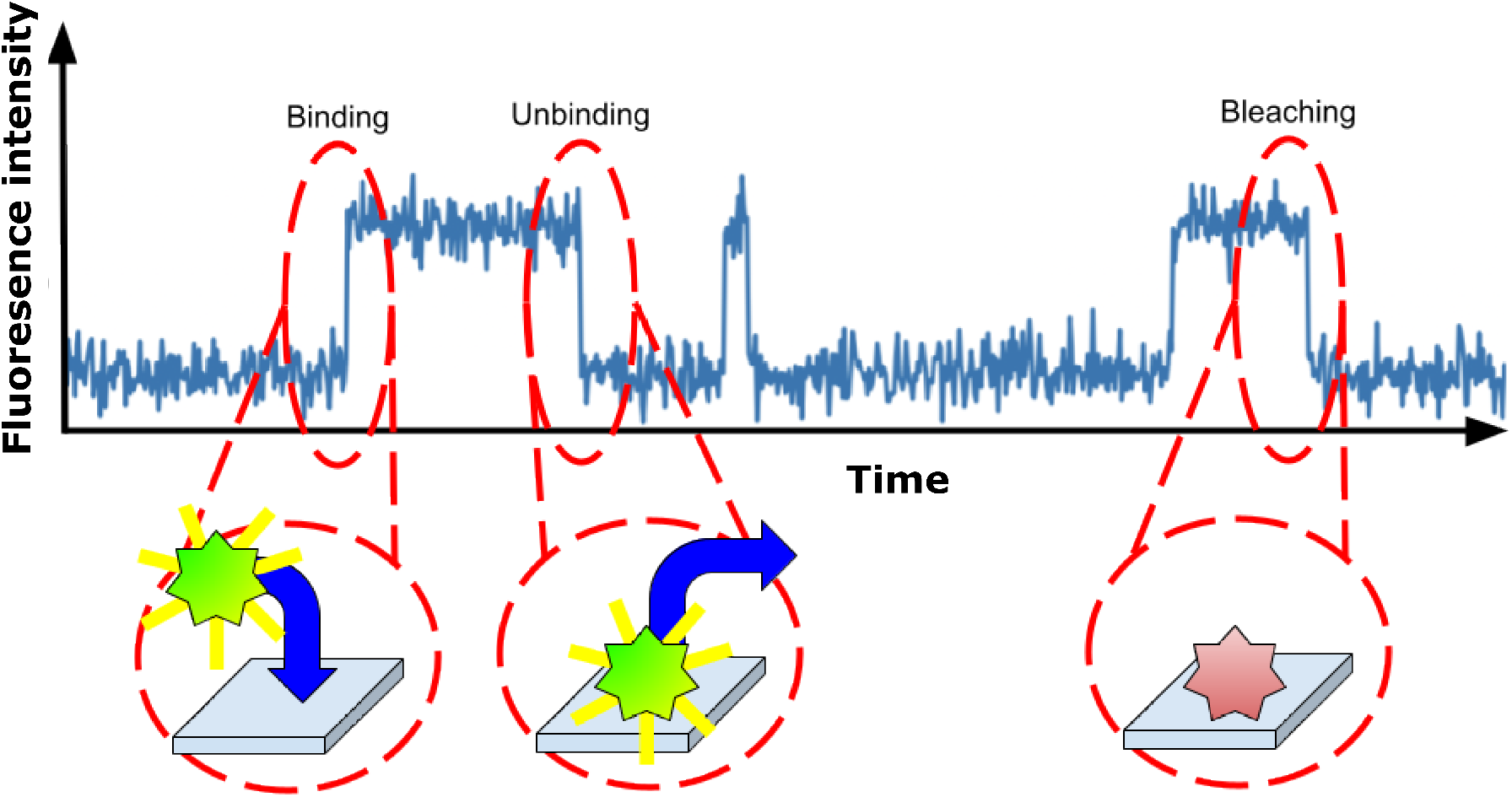
Illustration of a fluorescence intensity time trace. A hypothetical fluorescence intensity trace from a fluorescent ligand binding experiment. Circles are drawn around the three key regions indicating a binding event, an unbinding event, and a photobleaching event, respectively. Of note is the inability to distinguish unbinding from bleaching by eye.

Here, we propose a method to achieve accurate binding rates while considering photobleaching. We focus on inferring rates instead of transition probabilities because rates allow us to disentangle the kinetic contributions of binding from bleaching [45, 46]. Concretely, to model the stoichiometry of assemblies, we assume two photophysical states (bright and photobleached) and two binding states (bound and unbound) for each monomer. An illustration of our system can be found in figure 1. Our method determines the stoichiometry of assemblies by inferring binding rates alongside the photobleaching rates from intensity fluctuations over time. To achieve this, our method analyzes the sum of intensities of pixels around individual regions of interest (ROI) to infer state trajectory of the system, including both photobleaching and binding states for each ROI alongside all the rates. Such intensity traces are often given as a convolution of noisy measurement of photon counts over time due to photon shot noise and noises intrinsic to the detector devices such as readout noise [47]. Moreover, the kinetics of binding and fluorophore photobleaching are inherently random processes leading to additional layers of noise. We are able to separate the unbinding rates from photobleaching rates by utilizing information about the laser intensity of each trace. As such, we adopt a Bayesian framework to account for all the sources of uncertainty inherent to the problem. In particular, we leverage the Hidden Markov model (HMM) within a Bayesian framework to deal with the input noisy intensity traces [48, 49, 44, 50].

For the sake of computational efficiency, our framework consists of two parts or modules: in the first part, we learn the state trajectories using the noisy input data; in the second part, we use the inferred state trajectories to learn the rates. An important advantage of this two-step framework is that it allows us to simultaneously analyze different ROIs with different intensity statistics, drastically speeding up computation through parallelization. In this work, we illustrate the two-stage mathematical framework of our method and construct the likelihood and posterior. After which, we focus on the inverse model and benchmark our method using synthetic data, and *in vitro* experimental data obtained from DNA origami experiments.

## 2 Methods

### 2.1 Computational Methods

Our computational method is divided into two modules: State Inference and Rate Inference. In State Inference, for each individual ROI we determine how many fluorophores are present in the bright state at each time point given the intensity trace of the ROI. We then consolidate and pass these results to our Rate Inference step, where we use the state trajectories to infer the binding rates and photobleaching rates for the system. While in principle we could combine these two steps into a single analysis, inputting raw intensity traces to output rates, we break them up to drastically speed up computation via parallelization. The loss in accuracy due to splitting the analysis into two stages will be minimal, so long as data are in a regime where the brightness steps are clear in the intensity traces. On the other hand, if the SNR in the data is too low, then our analysis may fail, for example, if the standard deviation of the noise is much larger than the size of a step.

#### 2.1.1 State Inference

In the State Inference step of our analysis, we take in an intensity trace and output the state trajectory, which is the number of bright fluorophores at each time point. We define ***x***_1:*N*_, the “ brightness trace”, to be the ROI’s intensity at each time level with *x*_*n*_ the intensity at time *n*. We also define ***b***_1:*N*_, which we will refer to as the “ state trace”, to represent the number of bright fluorophores across the time points of the measurement where *b*_*n*_ is the number of bright fluorophores in the ROI at time *n*. Our goal in the State inference step is to find the most probable state trace, ***b***_1:*N*_, given a brightness trace, ***x***_1:*N*_.

In order to find the most probable state trace, ***b***_1:*N*_, given a brightness trace, ***x***_1:*N*_, we use Bayes’ theorem

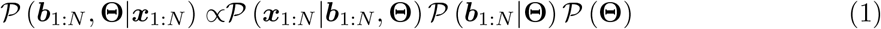

where **Θ** are model parameters and we split the posterior, *𝒫* (***b***_1:*N*_, **Θ**|***x***_1:*N*_), into a likelihood, *𝒫* (***x***_1:*N*_ |***b***_1:*N*_, **Θ**), a prior over brightness trace, *𝒫* (***b***_1:*N*_ **Θ**), and a prior over model parameters defined shortly, *𝒫* (**Θ**). For the likelihood, we assume each measurement independent and normally distributed around a mean given by the background plus the number of bright fluorophores,

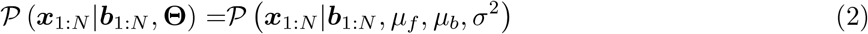

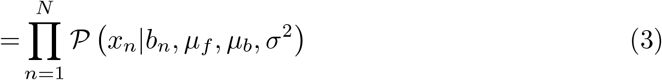

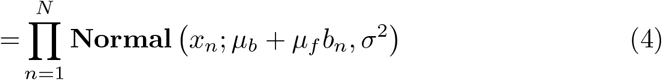

where we have introduced three variables: the fluorophore brightness, *µ*_*f*_, the background brightness, *µ*_*b*_, and the noise *σ*^2^. We will model the prior over fluorescence intensity trace as a Hidden Markov Model in which each state is categorically distributed according to the state at the previous time level [44],

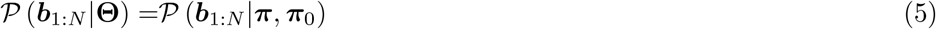

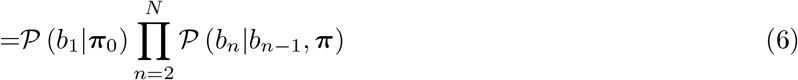

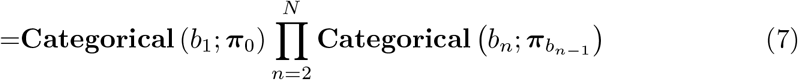

where we have introduced two variables: the state transition matrix, ***π***, and an array of starting probabilities ***π***_0_. The quantity 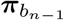 is interpreted as the “ row of ***π*** corresponding to the state at time level *n −* 1”. Each element, *π*_*ij*_, of ***π*** represents the probability that a time step will have *j* fluorophores given that the previous time step had *i* fluorophores. While in principle, there could be an unlimited number of fluorophores in the ROI at a given time step, in practice we will cap the possible number of fluorophores to *M*, meaning that ***π*** is a *M* + 1 by *M* + 1 matrix (since we include 0 fluorophores as a possibility). The full set of parameters can now be defined to include **Θ** = {*µ*_*f*_, *µ*_*b*_, *σ*^2^, ***π***_0_, ***π***}.

Working in the Bayesian paradigm, we must assign priors over the variables: *µ*_*f*_, *µ*_*b*_, *σ*^2^, ***π***_0_, and ***π***. We choose a Gaussian over *µ*_*f*_ and *µ*_*b*_, an inverse gamma for *σ*^2^, and Dirichlet distributions for ***π***_0_ and each row of ***π***, since they are the suitable conjugate priors of their respective likelihood distributions [44],

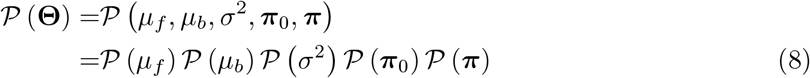

where the individual priors are given by

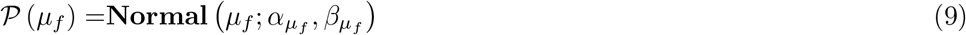

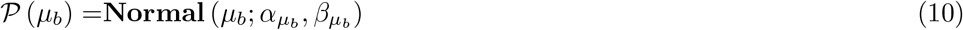

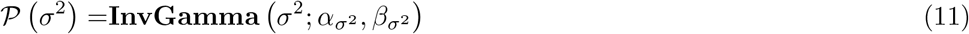

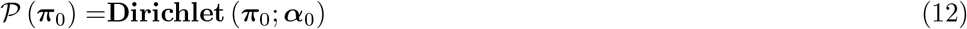

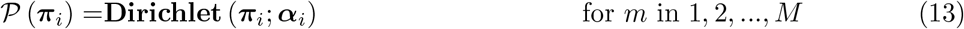

and where the *α*’s and *β*’s are hyperparameters. Altogether, the posterior over ***b***_1:*N*_ and **Θ** now reads

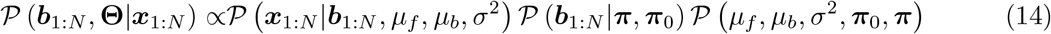

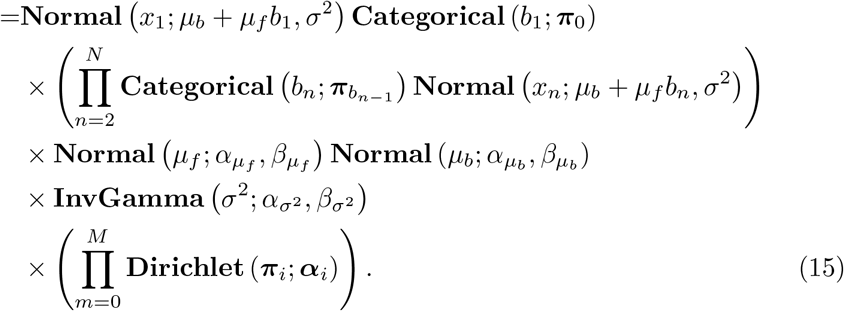

In the top of figure 2, we present our posterior as a graphical model [44]. In the graphical model, nodes (circles) represent random variables, and arrows represent conditional dependence.

**Figure 2:**
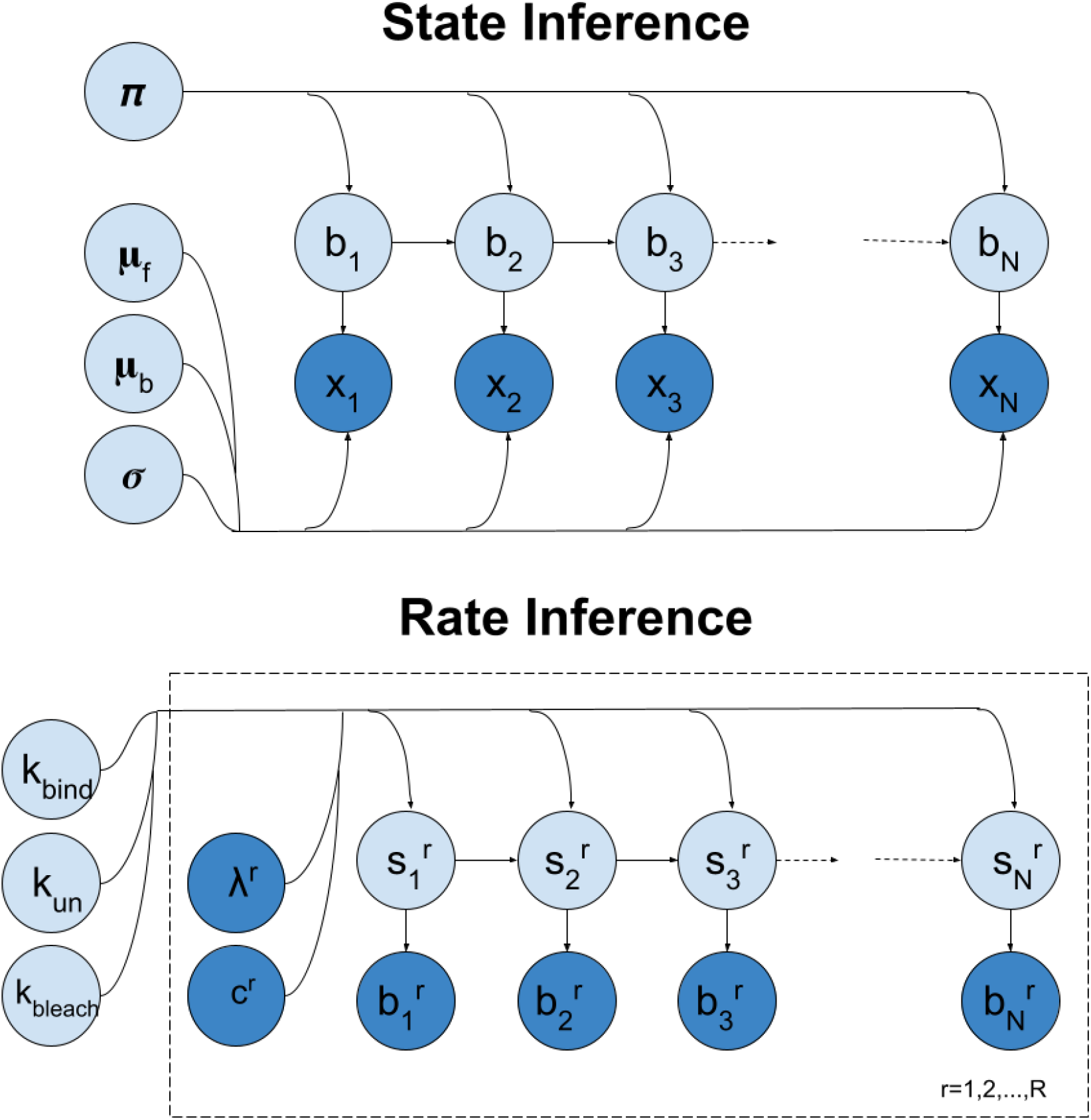
Graphical model of our analysis. Nodes (circles) represent random variables. Dark nodes represent measurements. Arrows between nodes represent conditional dependence. Plates (dashed rectangles) around nodes represent groupings of variables, which are repeated along the index in the lower right corner. In the top of this figure we present our graphical model for State Inference. In the bottom of this figure we present our graphical model for Rate Inference. All variables are defined in the main body.

We wish to find the value of ***b***_1:*N*_ that maximize equation (15), which we will refer to as the maximum a posteriori (MAP) sample. We will do so using Gibbs sampling [44], where we start with an initial guess then iteratively propose new values for variables, accepting and rejecting according to a ratio of the probabilities. We use Gibbs sampling, as opposed to optimization, to avoid local minima in our posterior. The algorithm is as follows:

- Start with initial set of variables, 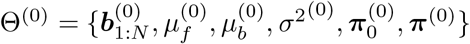.
- Set the MAP sample to this value, Θ_*MAP*_ = Θ^(0)^.
- For many iterations *i*:
  - Propose a new set of parameters Θ^*′*^ based on a proposal distribution *q*(Θ^*′*^|Θ^(*i−*1)^).
  - Calculate the acceptance ratio *a*, defined as:

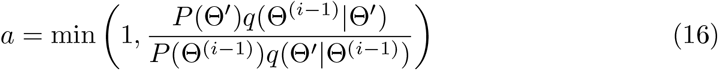

where *P* (Θ) represents the posterior probability of the parameters and *q*(Θ_*a*_| Θ_*b*_) represents the proposal distribution.
  - Accept or reject the new parameter set Θ^*′*^ with probability *a*.
  - Compare Θ^(*i*)^ to Θ_*MAP*_. If *P* (Θ^(*i*)^) *> P* (Θ_*MAP*_), update Θ_*MAP*_ = Θ^(*i*)^.
- The final Θ_*MAP*_ is the estimate for the maximum a posteriori value.

The final 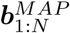 is the state trace. All the state traces are then fed into the next step, Rate Inference.

#### 2.1.2 Rate Inference

Once we have state traces for each ROI, we are ready to estimate the binding and photobleaching rates governing transitions. Here we assume there are *R* ROIs, with state traces 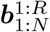 where 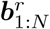 is the state trace of ROI *r* and 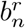 is the number of fluorophores in ROI *r* at time level *n*. We wish to infer the binding rate, *k*_bind_, the unbinding rate, *k*_un_, and the photobleaching rate, *k*_bleach_ given the state traces, 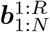, as well as the laser intensity of each trace, ***λ***^1:*R*^, and the concentration of binding agent at each trace, ***c***^1:*R*^.

We again invoke Bayes’ theorem to decompose the probability of our target rates, *k*_bind_, *k*_un_, and *k*_bleach_, given the state traces, 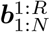 into a likelihood and prior

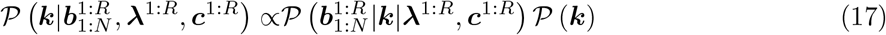

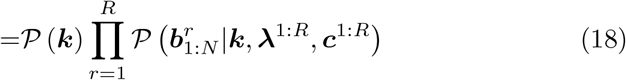

where ***k*** is the grouping of all rates (*k*_bind_, *k*_un_, and *k*_bleach_) and we have assumed that the likelihoods governing different ROIs are independent.

We now determine the likelihood of an ROI’s state trace 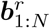. The state at each time point, 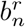, represents the number of fluorophores that are bright at the time level *n*. However, there are many different configurations of fluorophore binding and photophysics that can give rise to a single 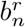. For example, even if we see only one bright fluorophore there could be other fluorophores bound in a photobleached state. We call 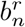 a “ macrostate” and denote all fluorophore configurations, 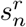, a “ microstate”, where a fluorophore configuration is the number of bright and photobleached fluorophores bound at the time level. We let Bright 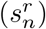 and Bleached 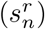 be two functions selecting the number of bright and photobleached fluorophores in the microstate. We can then enumerate the possible microstates as follows

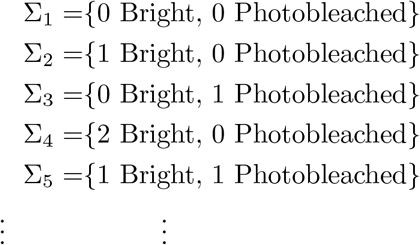

where Σ_*i*_ is the *i*th macrostate, and we enumerate up until we exhaust all *M* available fluorophores. The likelihood of a state trace, 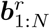 is the probability of a microstate trace that gives rise to the observed macrostate trace, marginalized over all possible microstate traces. Assuming that microstates follow a Hidden Markov Model this becomes,

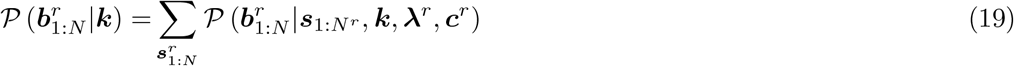

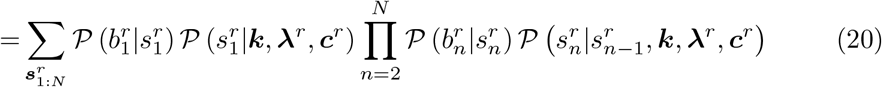

where 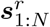 represents the full microstate trace. In the bottom of figure 2, we present our posterior as a graphical model [44]. In the graphical model, nodes (circles) represent random variables, arrows represent conditional dependence, and the plate (rectangle) represents groupings of random variables repeated over an index.

Equation (20) requires a form for the probability of microstate given a macrostate. The probability of the microstate given a macrostate is simply an indicator function, or delta function, equal to 1 when the number of bright fluorophores in the microstate is equal to the macrostate and 0 otherwise

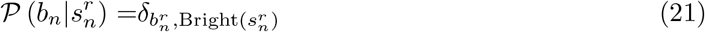

where *δ*_*ij*_ is the Kronecker delta. The microstate transition probability is calculated using the transition rates, *k*_bind_, *k*_un_, *k*_bleach_. Assuming one event per time level per ROI there are only four possible events: no transition, bright fluorophore binding, bright fluorophore unbinding, and photobleached fluorophore unbinding. We exclude the possibility of photobleached fluorophores binding, which is assumed to be rare. Letting Bright 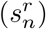 and Bleached 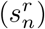 be functions which count the number of bright and photobleached fluorophores in state 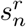 respectively, we can calculate the probability of each event as follows

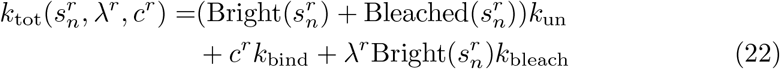

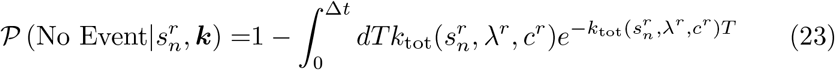

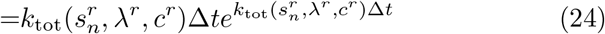

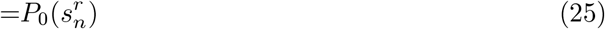

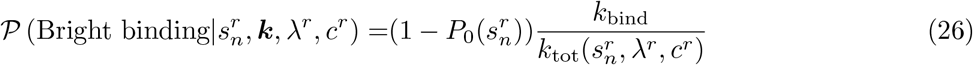

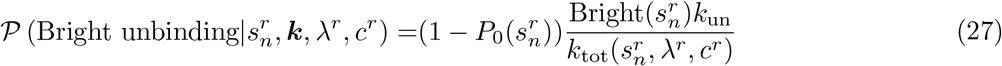

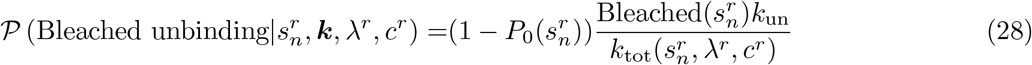

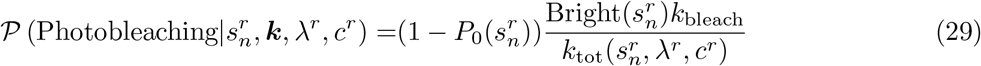

where 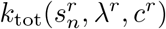 is the total rate, 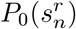 is the probability of a self transition (calculated as 1 minus the probability of an exponentially distributed event), and the remaining events are calculated according to their relative rates [51]. We note that the total binding rate is assumed to scale linearly with the concentration and the total photobleaching rate is assumed to scale linearly with laser intensity. These rates can be used to construct a master transition probability matrix, **Π**(*k*_bind_, *k*_un_, *k*_bleach_), where each element can be calculated by looking at changes in state populations between the initial state and final state for the 5 possible transitions: 1) No transition, 2) Binding, 3) Bright Unbinding, 4) Bleached unbinding, and 5) Photobleaching

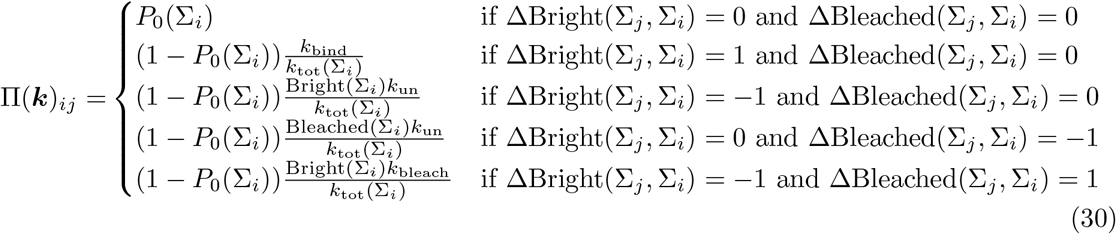

where

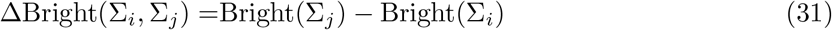

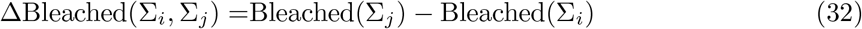

calculate the change in bright and bleached populations. We use the matrix presented in equation (30) to marginalize over the trajectories in equation (20) using the Forward-Backward algorithm [52], thus our likelihood, 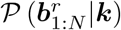, can be calculated exactly [52].

In order to find the set of rates, ***k***, that maximizes the posterior, equation (18), we again use a Gibbs sampling scheme, similar to that laid out in the previous section, but this time saving all samples, not just the MAP. The algorithm is as follows,

- Start with initial set of variables, 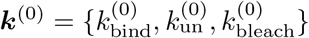
- For many iterations *i*:
  - Propose a new set of parameters ***k***^*′*^ based on a proposal distribution *q*(***k***^*′*^|***k***^(*i−*1)^).
  - Calculate the acceptance ratio *a*, defined as:

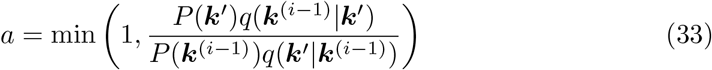

where *P* (***k***) represents the posterior probability of the parameters and *q*(Θ_*a*_ |Θ_*b*_) represents the proposal distribution.
  - Accept or reject the new parameter set ***k***^*′*^ with probability *a*.
  - Save the sample, ***k***^(*i*)^.

After drawing many samples, we may histogram the results to visualize the mean and variance of the distribution.

### 2.2 Experimental Methods

#### 2.2.1 DNA Origami

In order to test our method of inferring binding rates for different numbers of docking sites within a ROI, we designed a DNA Origami with varying numbers of external docking sites featuring the S1 sequence (see figure 3 and table SI1). The docking sites were added to the 3’ s-ends of selected staple strands positioned at 10 nm distances. For colocalization, the new rectangular origami (NRO) was equipped with three S2 docking sites (21 nt) for permanent external labeling of the structure with a Atto542-labeled complementary S2 strand (see table SI2). For immobilization of the DNA origami to a glass surface functionalized with Biotin-Streptavidin, six staple strands of the NRO were labeled with biotin on their 3’-end. Exact strand sequence can be found in the SI in table SI1, table SI2, and table SI3.

**Figure 3:**
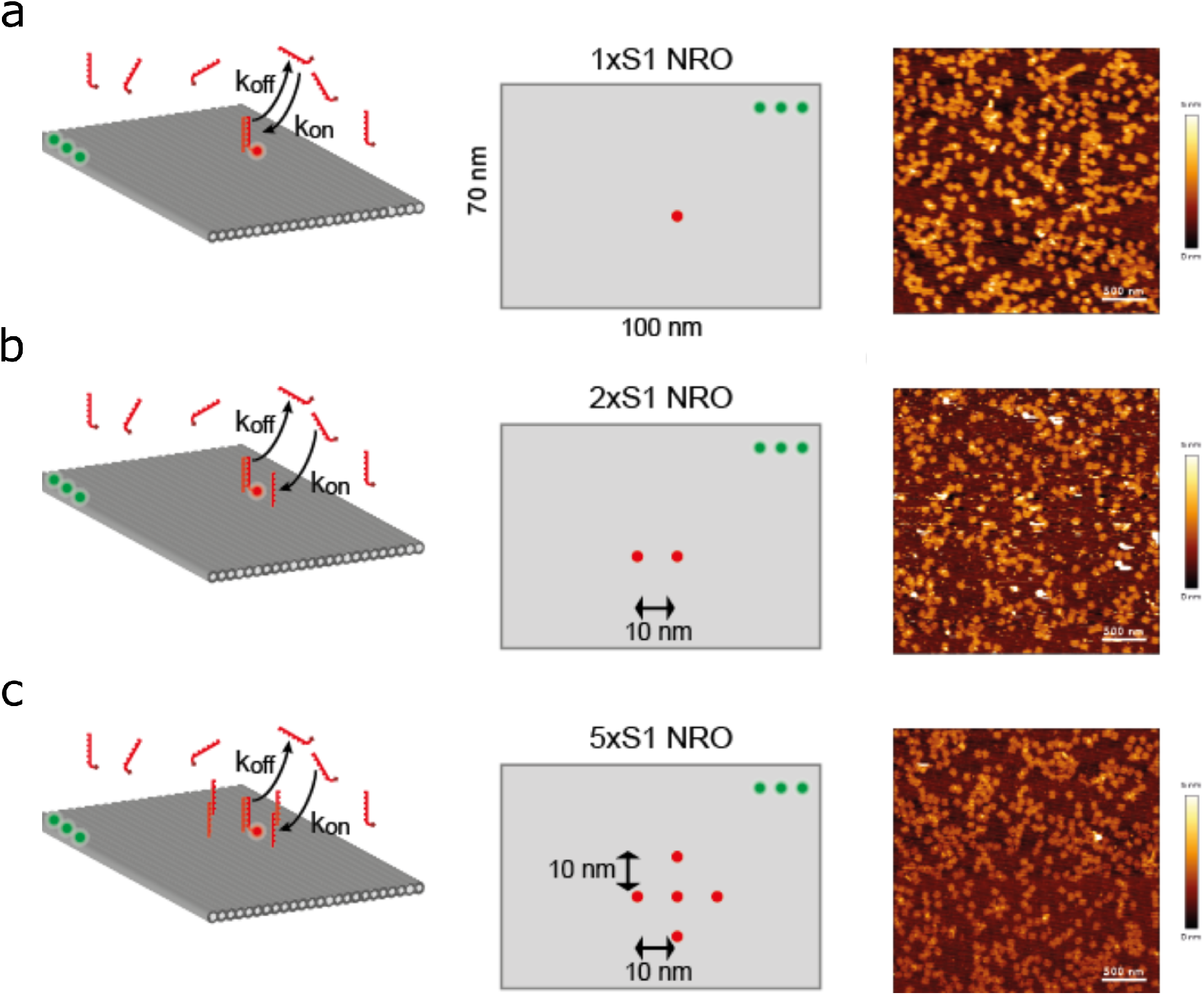
Illustration of the new rectangular DNA origami (NRO) structure and its binding site arrangement. Each row provides information about DNA origami with a set number of S1 docking sites-(a) one binding site, (b) two binding sites, (c) five binding sites. The left column shows cartoon of the DNA origami with the imager strands (red) binding to the S1 docking sites and a stable reference label on three S2 docking sites (green). The middle column shows a diagram of the layout of binding sites and reference labels on the DNA origami. The right column shows atomic force microscopy images of the DNA origami immobilised on a mica surface.

The DNA origami was folded using 20 nM of the scaffold p7249, 200 nM of each unmodified staple strand and 600 nM of each modified staple strand (biotinylated and external S1 labeling strands) in buffer containing 10 mM Tris, 1 mM EDTA at pH 8 and 12.5 mM MgCl_2_. The mixture was heated to 70 °C and then cooled down to 25 °C with a linear thermal annealing ramp of 1°C/min [53].

Sample purification was performed by filtration using Amicon Ultra filters (100 kDa molecular weight cut-off, Merck, Germany). The filter was first centrifuged with folding buffer for 7 min at 6000 g. The sample solution was then loaded into the filter and centrifuged for 15 min at 6000 g. Then 300 µL of folding buffer was loaded into the filter and centrifuged for 15 min at 6000 g, which was repeated once. After the three washing steps, the filter was inverted and placed into a new collection tube. The purified sample could then be collected by centrifugation for 2 min at 1000 g. The successful folding of structures was confirmed by AFM analysis. DNA origami solutions were stored at −20 °C until used.

To determine correct folding, AFM scans in aqueous solution (AFM buffer = 40 mM Tris, 2 mM EDTA, 12.5 mM Mg(OAc)2·4 H2O) were performed on a NanoWizard® 3 ultra AFM (JPK Instruments AG) (see figure 3). For sample immobilization, a freshly cleaved mica surface (Quality V1, Plano GmbH) was incubated with 10 mM solution of NiCl_2_ for 3 min. The mica was washed three times with ultra-pure water to remove unbound Ni^2+^ ions and air-dried. The dried mica surface was incubated with 1 nM sample solution for 3 min and washed with AFM buffer three times. Measurements were performed in AC mode on a scan area of 3 x 3 *µ*m with a BioLeverMini cantilever (*ν*res = 110 kHz air / 25 kHz fluid, kspring = 0.1 N/m, Bruker AFM Probes).

#### 2.2.2 Sample preparation for fluorescence microscopy

Samples were prepared in chambered coverslips (LabTek) which were cleaned twice with 0.1 M HF with two water rinses after each cleaning step. The chambers were then incubated for 5 min in magnesium-free phosphate buffered saline (PBS). The coverslips were then coated with 1 mg/ml biotin-BSA overnight before being washed three times in PBS. Next, the coverslips were incubated for 30 min with 0.2 mg/ml streptavidin before they were washed three times with Red Base composed of 2x PBS with additional 0.5 M NaCl and 0.05% Tween20. 10 fmol of rectangular DNA origami was then incubated for 30 min on the surface of the coverslip, before removing the solution and washing three times with Red Base. Each DNA origami construct was pre-labeled with Atto542 as a reference marker and had a defined number of S1 docking sites (see Supplement). The samples were inspected for sufficient DNA origami density and in cases of low coverage additional DNA origami was added to achieve sufficient coverage. Each chamber of the coverslips was then filled with imaging buffer and encapsulated with self-sealing Parafilm(M). The imaging buffer contained 2.1 mM ascorbic acid (AA), 1 mM methyl viologen (MV), 2.5 mM protocatechuic acid (PCA) and 50 nM protocatechuate-3,4-dioxygenase (PCD) in Red Base. Different concentrations of Cy5-conjugated oligonucleotides were used in each experiment (1 nM, 2 nM, 5 nM, 10 nM). The imager strand concentration was adjusted by substituting Red Base with 1-10 µl of 1 µM imager strand diluted in Red Base.

#### 2.2.3 Data Acquisition

The samples were imaged on a custom microscope setup using an inverted microscope (Nikon Eclipse Ti, Nikon, Japan) and a fibre-coupled multilaser engine (MLE-LFA, TOPTICA Photonics, Germany). The excitation was coupled to the Nikon manual TIRF unit and then directed to 100×, 1.49 NA oil immersion objective (Apo TIRF, Nikon, Japan) via a four-band dichroic mirror (405, 488, 532, 640 nm). Emission was then filtered by 525/50 nm, 605/70 nm, and 690/70 nm bandpass filters (all AHF Analysetechnik, Germany) mounted on a motorized filter wheel (FW212C, Thorlabs, Germany) and collected by an EMCCD (Electron Multiplying Charge-Coupled Device) camera (iXon Ultra 897, Andor, UK) with a 105.6 nm pixel size. Initially, the fields of view were imaged for five frames in the 561 nm channel to attain reference data for localisations of the origami. Afterwards, the same positions are imaged at 640 nm under TIRF illumination with 21.7 mW and 43.8 mW measured before the objective with a power meter kit (PM120D, Thorlabs, Germany). These correspond to 98 W/cm^2^ and 198 W/cm^2^ at the focal plane, respectively. To gather kinetic data 8000 frames were recorded at 100 ms exposure (124.6 ms frame time) over a total duration of 16.5 min and saved as an image stack.

#### 2.2.4 Post Acquisition Data Processing

For each image stack, a custom Fiji script [54] determines the brightest values of each pixel, creating a maximum intensity projection, after which ThunderSTORM [55] is used to localise the coordinates for each region of interest (ROI), i.e. the location of DNA origami, in both channels. Then a Python script, which utilizes the trace extraction function of the quickPBSA library [37], is used to determine the intensity trace inside each ROI and subtract the background around that ROI. Then the maximum intensity projections of the two channels were registered using the MultiStackReg plugin [56] in Fiji giving a correction matrix used to transform the coordinates in the reference channel via a custom Fiji script. Finally, the ROIs further than 200 nm of the reference channel are considered unspecific binding and filtered out.

## 3 Results

The main goal of our algorithm is to learn the binding (*k*_bind_), unbinding (*k*_*un*_), photobleaching (*k*_bleach_) rates and the state trajectories given a set of noisy intensity traces 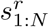 for every given ROI *r*. To do so, our algorithm takes a chain of numerical samples from the posterior using Monte Carlo methods as detailed in section 2. As such, our results are in the form of histograms of the numerical samples taken from the posterior for every individual unknown parameter, where the uncertainty of rate inference given a microstate trace over each parameter is reflected in the width of the corresponding histograms.

### 3.1 Simulated data

We first benchmarked our method using data simulated under different regimes data using the forward model described in the Methods (section 2). Starting with a base set of parameters chosen to mimic the experimental data, we ran inference while varying one parameter at a time to probe the robustness of our method with respect to different data paradigms. The base set of parameters chosen to qualitatively match the data is as follows: *k*_bind_ = 1 mHz/nM, *k*_un_ = 25 mHz, *k*_bleach_ = .01 mHz, *µ*_*f*_ = 400 ADU, *µ*_*B*_ = 0 ADU, *σ* = 2500 ADU, *dt* = 124 ms, and 2000 ROIs with traces simulated with either *λ* = 50 or *λ* = 100 in even proportion and *c* = 1 nM, *c* = 2 nM, or *c* = 5 nM in even proportion. We simulated data varying *k*_bind_, *k*_un_, *k*_bleach_, and the number of ROIs summarized in table SI4.

Initially, we probed the model’s robustness with respect to binding rate, *k*_bind_, by comparing analysis of data simulated with *k*_bind_ = 1 mHz/nM, *k*_bind_ = 2 mHz/nM, and *k*_bind_ = 5 mHz/nM. Figure 4 shows results from the data simulated with these different *k*_bind_ rates. Figure 4 is broken into three columns where the left column shows representative data traces, the middle column shows the inferred *k*_bind_ binding rate, and the right column shows the inferred *k*_un_ unbinding rate. For the plots showing data traces, we show two representative traces taken from the dataset, stacked on top of each other. For the plots showing inferred rates, the blue indicates the probability distribution (histogram of samples from the MCMC), and the red line indicates the ground truth value used in the simulation. As seen in the middle column of figure 4, our method accurately (within 10%) infers the correct *k*_bind_ rate with the analysis returning 1.031 ± 0.012, 1.971 ± 0.018 and 4.588 ± 0.030 mHz/nM for the *k*_bind_ value. *k*_un_ results remain also constant only slightly lower than the set 25 mHz.

**Figure 4:**
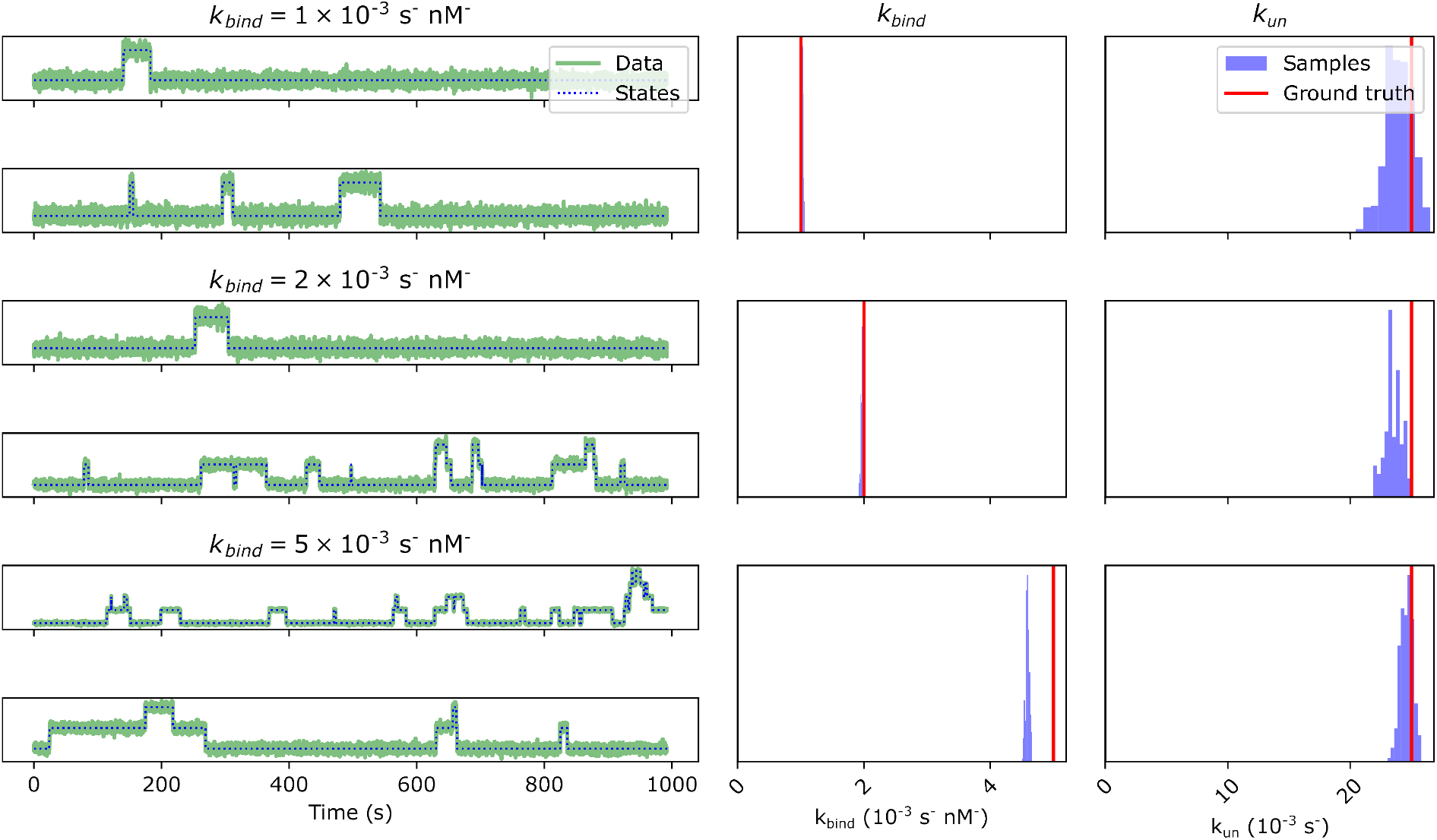
Simulated data with different *k*_bind_ binding rates. The top two rows of the left column show example traces from data simulated with *k*_bind_ = 1 mHz/nM, the middle two rows of the left column show example traces from data simulated with *k*_bind_ = 2 mHz/nM, and the bottom two rows of the left column show traces from data simulated with *k*_bind_ = 5 mHz/nM. The middle column shows inferred posteriors over binding rates, with blue indicating the probability distribution and a red line indicating the ground truth used for simulation. The right column shows inferred posteriors over unbinding rates, with blue indicating the probability distribution and red lines indicating ground truth values used for simulation. Dataset included 200 traces per condition. The parameters used in simulation are specified in main text and table SI4.

We next probed the model’s robustness with respect to unbinding rate, *k*_un_, by comparing analysis of data simulated with *k*_un_ = 10 mHz, *k*_un_ = 20 mHz, and *k*_un_ = 50 mHz. Figure 5 shows the results on data simulated with these different *k*_un_ rates. As seen in the far right column of figure 5, our model accurately infers the correctly the change of *k*_un_ rate with 9.623 ± 0.612 mHz, 23.950 ± 1.086 mHz, 45.650 ± 1.586 mHz for the 10 mHz, 20 mHz and 50 mHz, respectively. In line with the input, the *k*_bind_ results remained around within 0.1 of the 1 mHz/nM input for all simulations.

**Figure 5:**
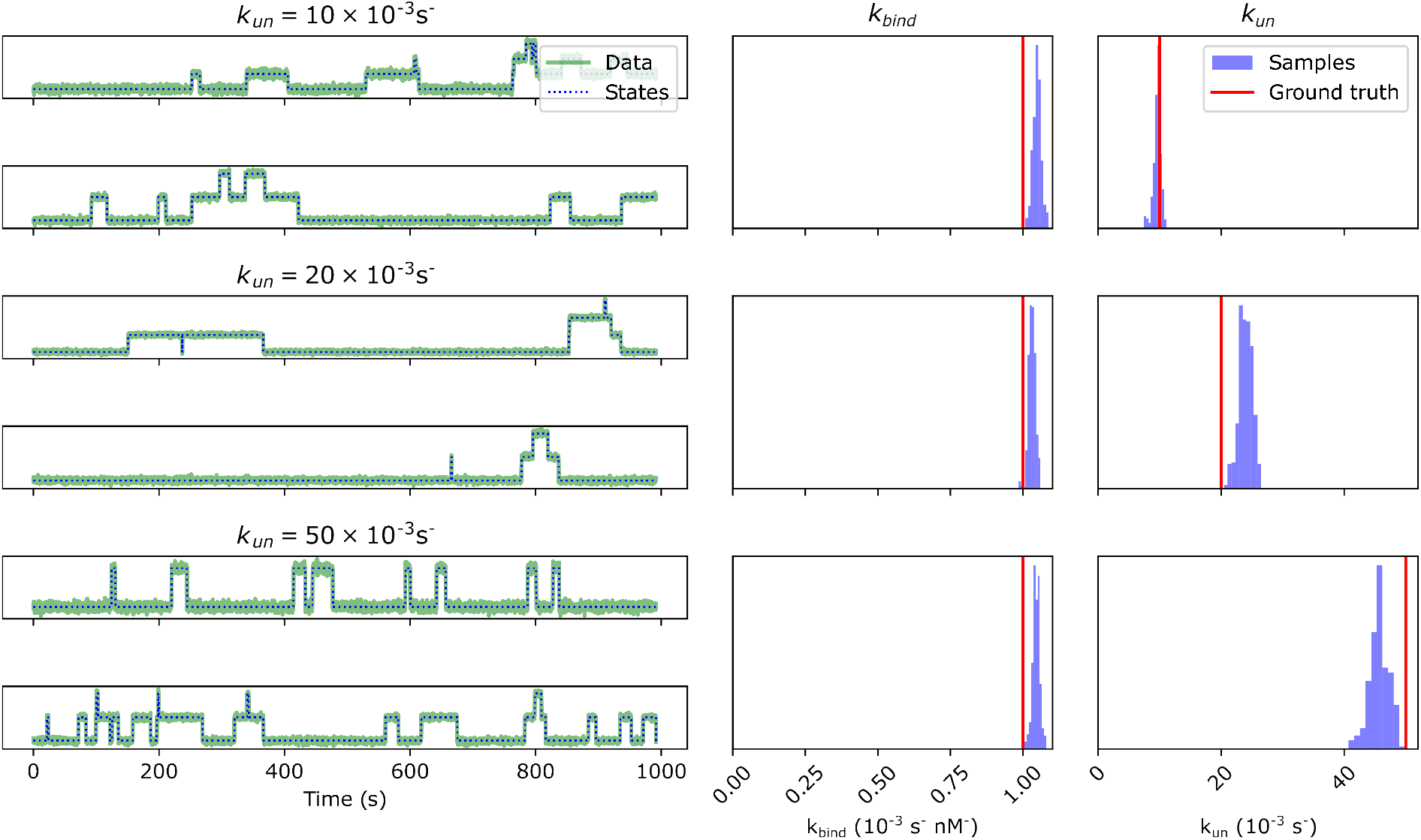
Simulated data with different *k*_un_ unbinding rates. The top two rows of the left column show example traces from data simulated with *k*_un_ = 10 mHz, the middle two rows of the left column show example traces from data simulated with *k*_un_ = 20 mHz, and the bottom two rows of the left column show traces from data simulated with *k*_un_ = 50 mHz. The middle column shows inferred posteriors over binding rates, with blue indicating the probability distribution and a red line indicating the ground truth used for simulation. The right column shows inferred posteriors over unbinding rates, with blue indicating the probability distribution and red lines indicating ground truth values used for simulation. Dataset included 200 traces per condition. The parameters used in simulation are specified in main text.

After that, we probed the model’s robustness with respect to photobleaching rate, *k*_bleach_, by comparing analysis of data simulated with *k*_bleach_ = 0.01 mHz, *k*_bleach_ = 0.02 mHz, and *k*_bleach_ = 0.05 mHz. Figure 6 shows the results on data simulated with these different *k*_bleach_ rates. As seen in figure 6, we are still able to infer accurate *k*_bind_ and *k*_un_ rates for all values of *k*_bleach_.

**Figure 6:**
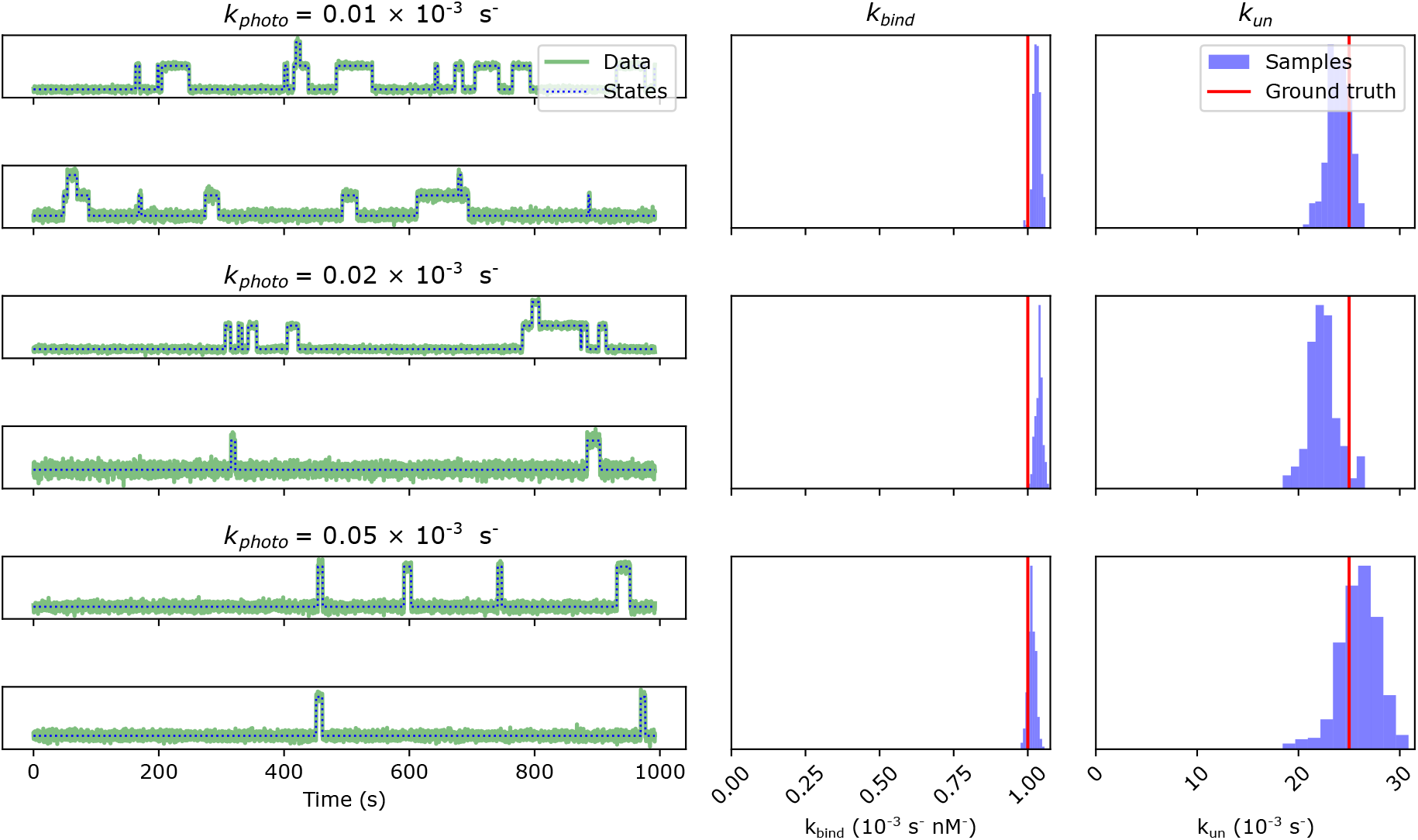
Simulated data with different *k*_bleach_ photobleaching rates. The top two rows of the left column show example traces from data simulated with *k*_bleach_ = .01 mHz, the middle two rows of the left column show example traces from data simulated with *k*_bleach_ = .02 mHz, and the bottom two rows of the left column show traces from data simulated with *k*_bleach_ = .05 mHz. The middle column shows inferred posteriors over binding rates, with blue indicating the probability distribution and a red line indicating the ground truth used for simulation. The right column shows inferred posteriors over unbinding rates, with blue indicating the probability distribution and red lines indicating ground truth values used for simulation. Dataset included 200 traces per condition. The parameters used in simulation are specified in main text and table SI4.

Finally, we probed the model’s robustness with respect to number of ROIs, by comparing data simulated with 2000 ROIs, 1000 ROIs, and 400 ROIs. Figure 7 shows the results on the datasets simulated with these different numbers of ROIs. Despite the small increase in uncertainty, marked by the spread in probability, our method was able to accurately learn *k*_bind_ and *k*_un_ from even a small amount of data.

**Figure 7:**
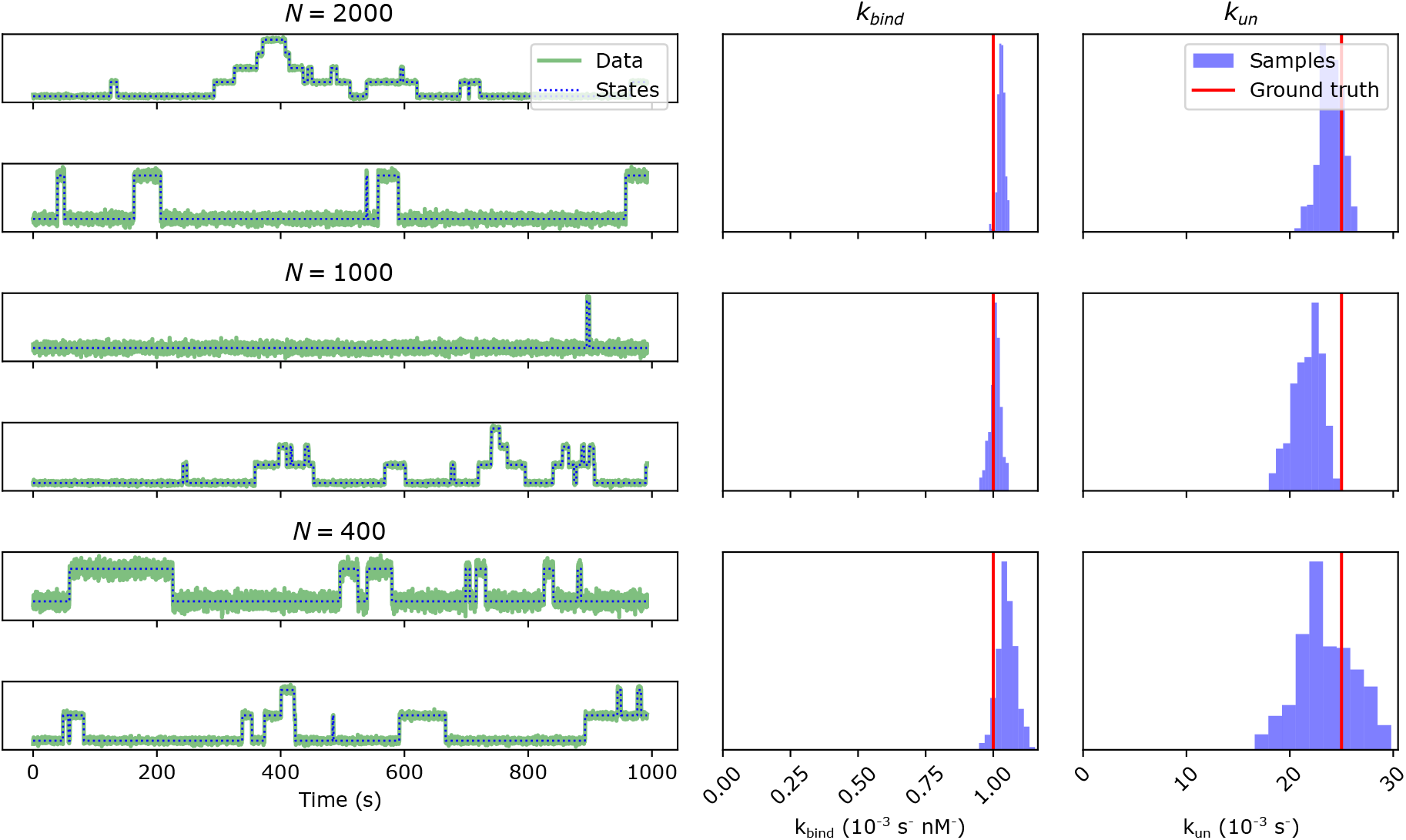
Simulated data with different number of ROIs. The top two rows of the left column show example traces from data simulated with 2000 ROIs, the middle two rows of the left column show example traces from data with 1000 ROIs, and the bottom two rows of the left column show traces from data with 400 ROIs. The middle column shows inferred posteriors over binding rates, with blue indicating the probability distribution and a red line indicating the ground truth used for simulation. The right column shows inferred posteriors over unbinding rates, with blue indicating the probability distribution and red lines indicating ground truth values used for simulation. The parameters used in simulation are specified in main text and table SI4.

### 3.2 Experimental data

We applied our method to data taken from highly controlled DNA origami experiments in which fluorescently labeled particles were allowed to bind and unbind to origami with 1, 2, and 5 binding sites. Results for inference on experimental data are shown in figure 8. Each row of figure 8 represents an experiment with a different number of binding sites with the inferred rates of binding and unbinding shown as a histogram.

**Figure 8:**
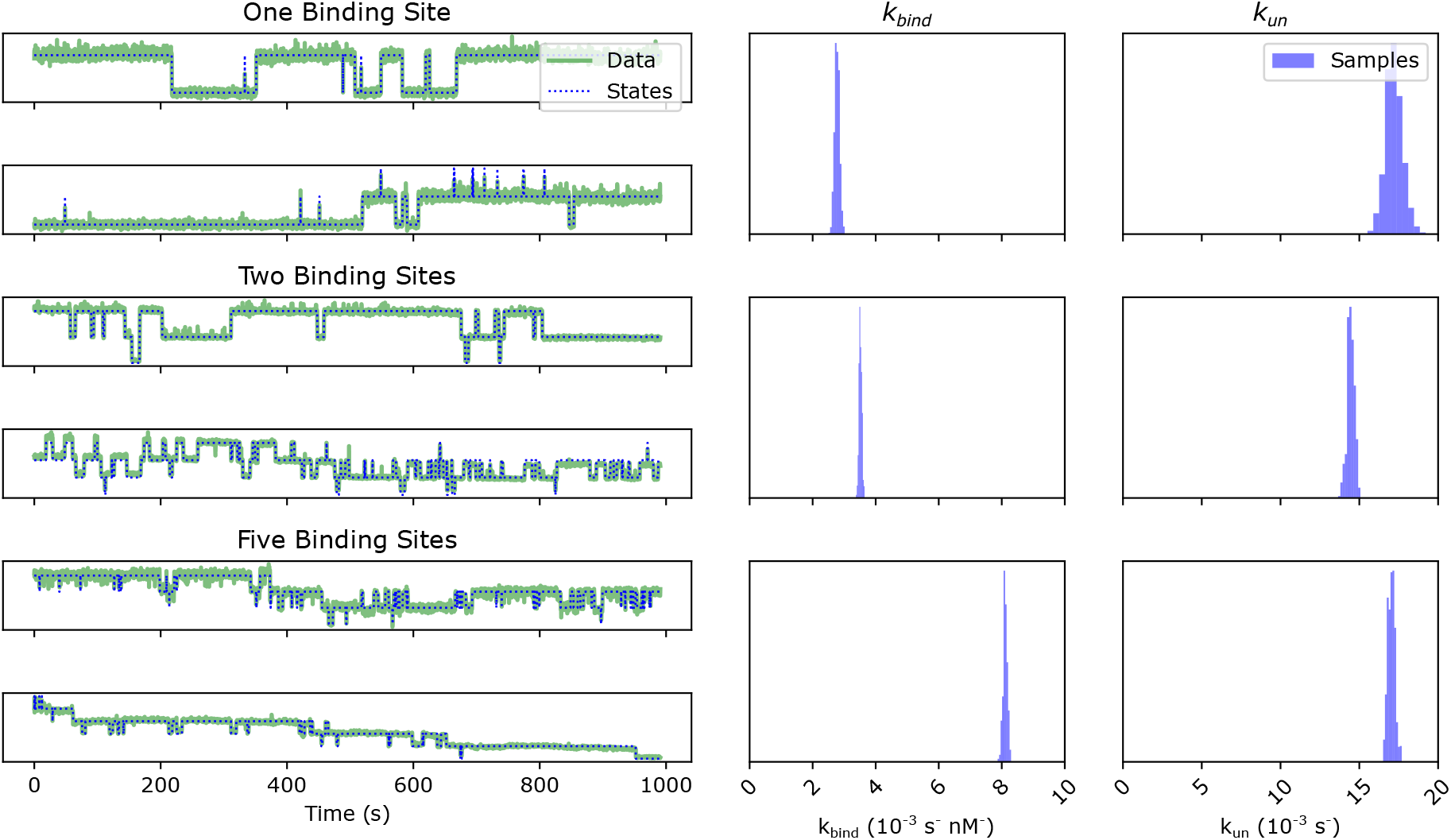
Data from DNA origami binding experiments. The left column shows example traces from our experiments. The top two rows of the left column show example traces from data with one binding site, the middle two rows of the left column show example traces from data with two binding sites, and the bottom two rows of the left column show traces from data with five binding sties. The middle column shows inferred posteriors over binding rates. The right column shows inferred posteriors over unbinding rates. Data from 126, 393 and 353 traces were analyzed for the one, two and five binding site samples, respectively.

We cannot compare our inferred rates directly to a ground truth, as the ground truth is unknown. However, we can evaluate the accuracy of our model by checking to see if the output behavior matches expectations. For example, the binding rate should scale with the number of available binding sites, but the unbinding rate should be unaffected by the number of binding sites. As seen in figure 8, inferred rates qualitatively increase with binding sites. Our inferred binding rates are the 2.77 *±* 0.07 mHz/nM (*mean±SD*) for 1 binding site, 3.52 *±* 0.04 mHz/nM for 2 binding sites, and 8.1 *±* 0.07 mHz for 5 binding sites, which increases with concentration. These values also match the values seen in literature when it comes to a 10 nucleotide imager strands [57]. Meanwhile, the unbinding rates are 17.2 *±* 0.5 mHz, 14.4 *±* 0.2 mHz, and 17.0*±* 0.2 mHz, effectively remaining constant. The experimental data is intrinsically noisy, containing many artifacts, such as spikes (see figure 8), which complicate analysis. These can be attributed to unbound strands moving over the ROI or the molecule escaping a prolonged dark state. The bleaching rates show a small values not correlated to the binding site number (see table SI5).

## 4 Discussion

Quantifying binding kinetics while overcoming noise and photobleaching artifacts is important as it provides a deeper and more accurate understanding of molecular interactions and behaviors in chemical and biological systems. This enhanced understanding is crucial in dissecting the intricacies of protein self-assembly [1, 2, 3], enzymatic and protein-protein interactions [5, 6, 7, 8, 9], DNA-protein interactions [11, 12, 13], and broader cellular processes, such as nuclear dynamics during mitosis and [15, 16, 17].

Here, we presented a Bayesian inference scheme for inferring binding rates separately from photobleaching rates, by directly monitoring fluorescence intensity over time. We prioritized inferring rates over transition probabilities, as it enables us to disentangle the contributions of binding and photobleaching. Our method is broken into two modules: a State Inference algorithm that infers the number of fluorophores at each time level directly from the data, and secondly a Rate Inference step which infers the binding rates from the output fluorescence trace. We benchmarked our method on simulated data, inferring reliable results. We showed that when run on experimental data we inferred expected behavior of increasing binding rates with constant unbinding rates for increasing concentrations.

A significant way to enhance this work would be by incorporating spatial information. Currently, the method works by analyzing intensity traces from isolated ROIs. By incorporating spatial information we would be able to take into account other potential noise generating artifacts such as uneven illumination of ROIs [58].

Another way to improve this work would be to incorporate additional photophysical states. For example, in the case of photo-blinking the number of brightness steps is not equal to the number of fluorophores, making analysis with traditional HMMs impossible [39]. Our method, on the other hand, is generalizable to an arbitrary number of dark and bright states. However, while a state inference analysis can be effective when state traces are constrained, as in the case of counting by photobleaching where the fluorophores are assumed to start bound and end bleached with no unbinding [39], without such a constraint the number of possible state trajectories becomes combinatorically larger. As a result it may be difficult, for fundamental reasons, to perform scalable analysis with a higher number of photophysical states.

## Data Availability

Data used in this work can be found at labpresse.com.

## Code Availability

Code used in this work be found at labpresse.com.

## Acknowledgements

SP acknowledge support from an NIH MIRA award (NIGMS R35GM148237). DPH acknowledges the support from the Centre of Membrane Proteins and Receptors (COMPARE) funded by the Universities of Nottingham and Birmingham as well as support from the Academy of Medical Sciences (Grant APR2*\*1013).

## Author Contributions

SP and DPH conceived of the project. MS designed and created the DNA origami. SAT performed the sample preparation, imaging and trace extraction for the experimental data. JSB carried out the coding and development. JSB and MF performed the analyses and wrote the manuscript.

## Competing Interests

SP and JSB acknowledge a competing interest with their affiliation with Saguaro Solutions.

## Supplementary Information

**Table SI1:**
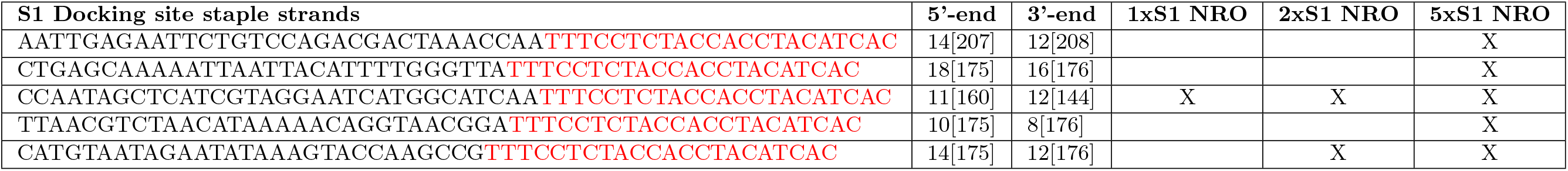
External S1 docking site staple strands of the NRO DNA Origami. Sequences are denoted from 5’-to 3’-end. The S1 docking site staple strands exhibit an over 20nt long docking site, marked in red, on the 3’-end. The numbers for the 5’-end 3’-end of the staples represent the helix number in the corresponding caDNAno file. Number in brackets represent the starting and ending position of the staple in the corresponding helix.

**Table SI2:**
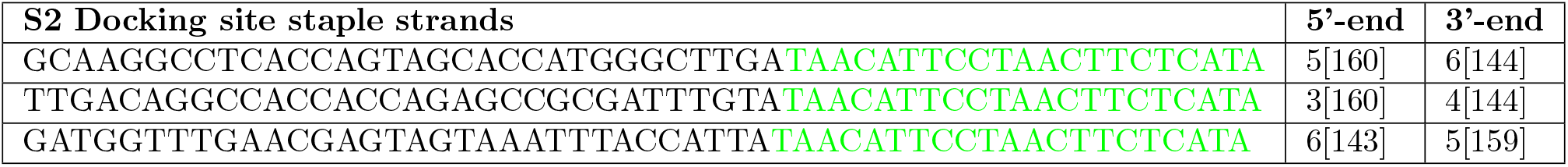
Modified staple strands of the NRO DNA Origami. Sequences are denoted from 5’-to 3’-end. The three S2 docking site staple strands exhibit an over 20nt long docking site, marked in green for colocalization, on the 3’-end. For immobilization, the biotinylated staple strands are modified with biotin on the 3’-end. The numbers for the 5’-end 3’-end of the staples represent the helix number in the corresponding caDNAno file. Number in brackets represent the starting and ending position of the staple in the corresponding helix.

**Table SI3:**
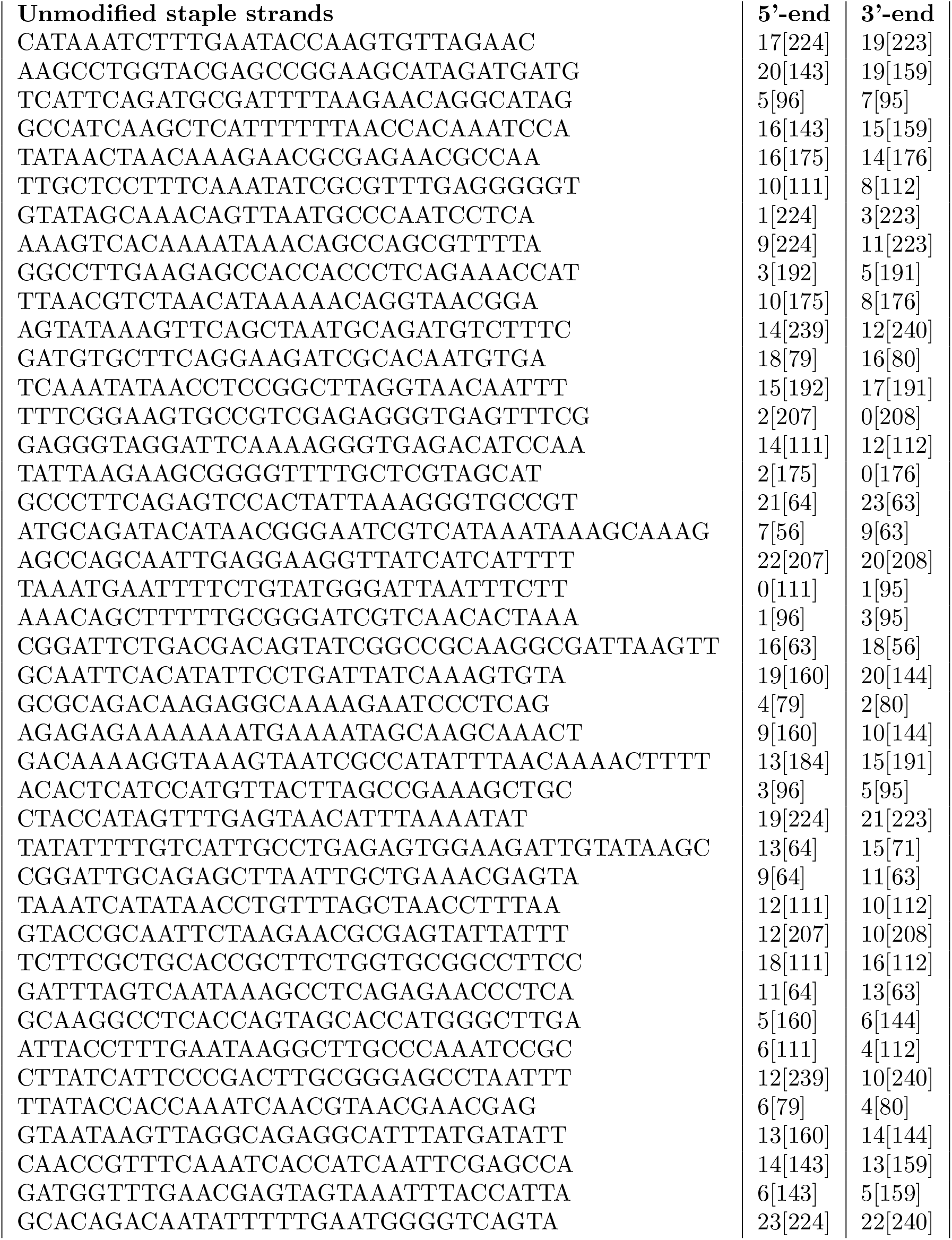
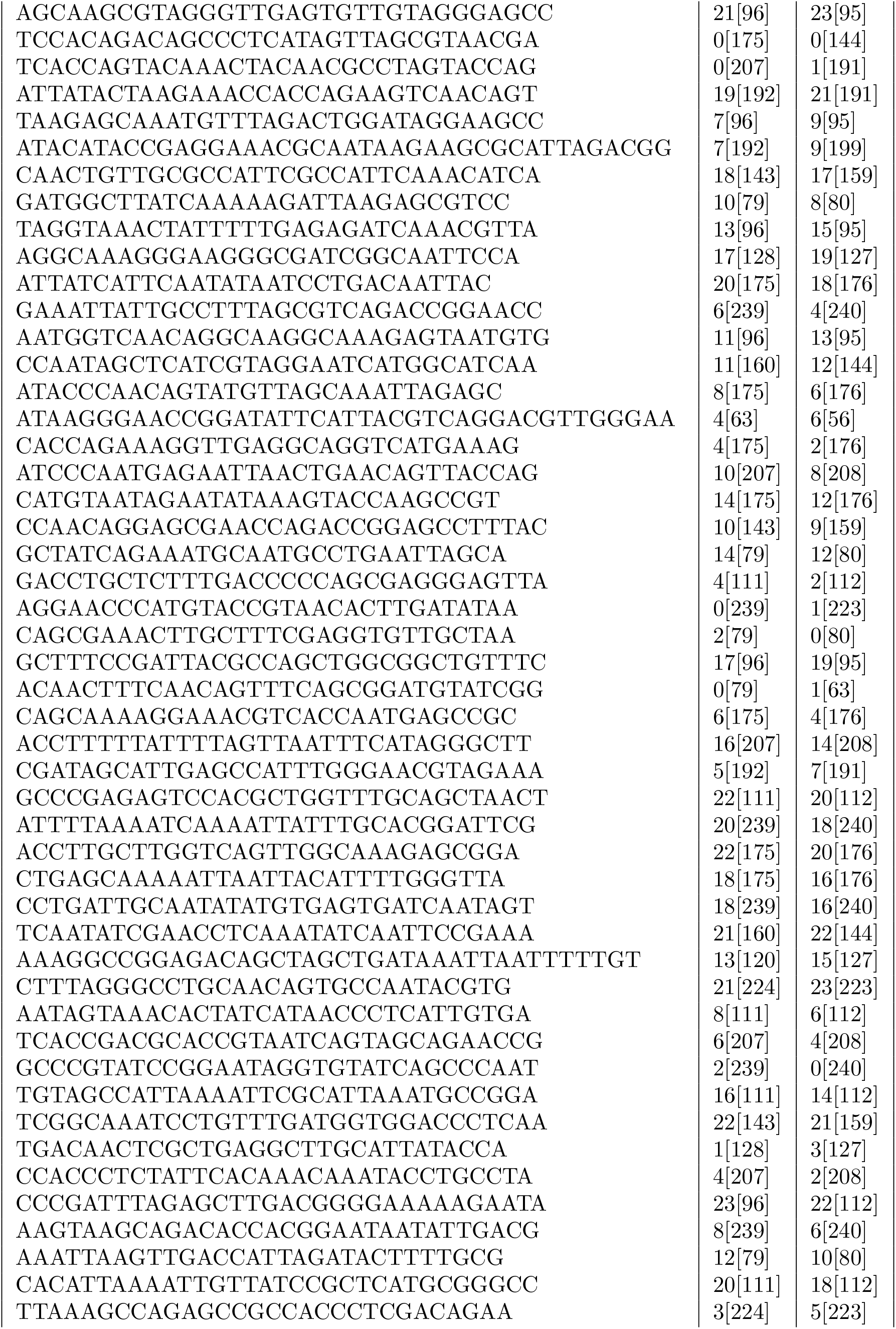
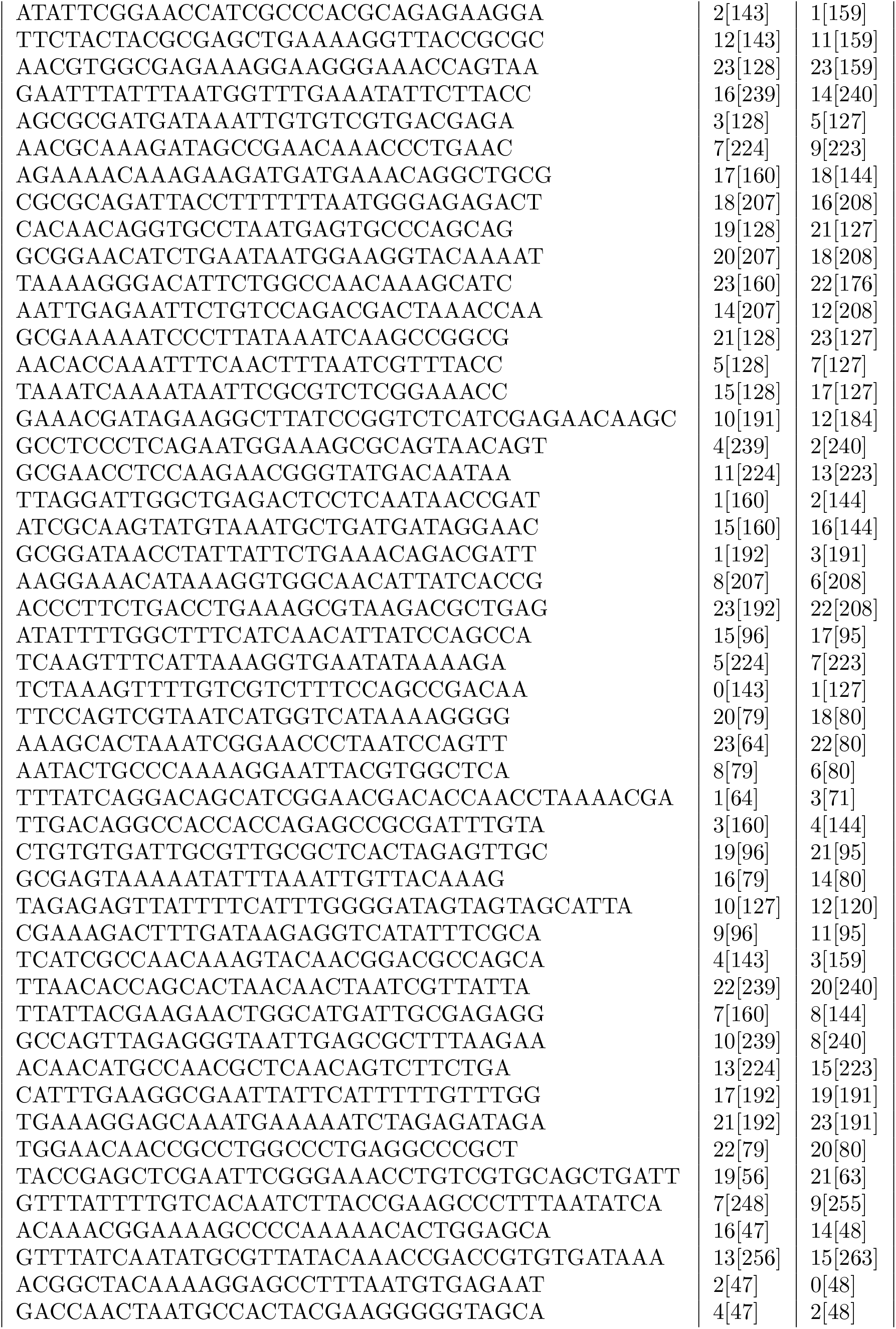
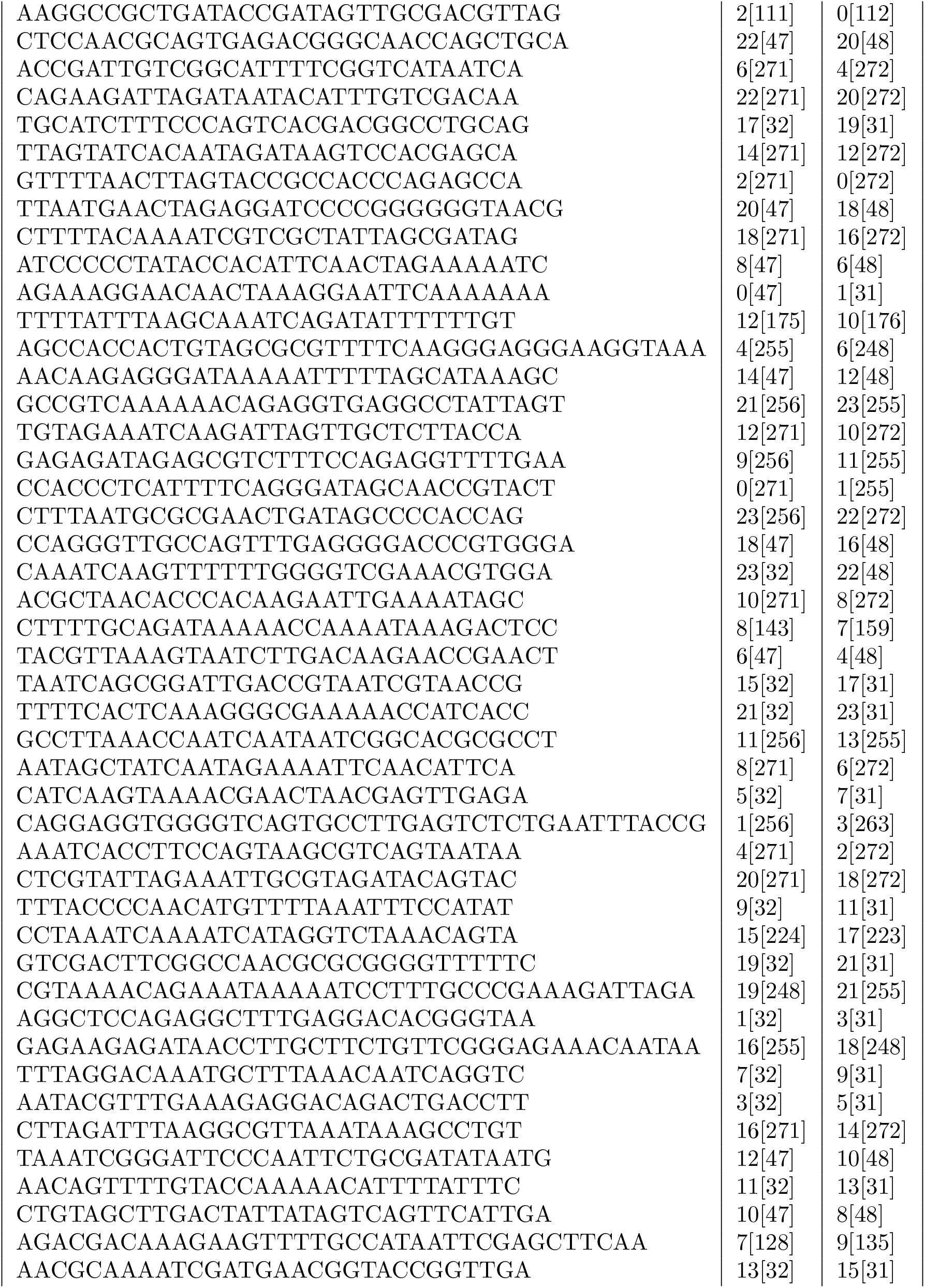
Unmodified staple strands of NRO DNA origami. Sequences are denoted from 5’-to 3’-end. The numbers for the 5’-end 3’-end of the staples represent the helix number in the corresponding caDNAno file. Number in brackets represent the starting and ending position of the staple in the corresponding helix.

**Table SI4:** Parameters used to create the simulated data. The three values were used in the forward model described in section 2. All data for figure 7 shares the same parameters with variation only in the number of data points.

| Figure | $k_{\text{bind}}(s^{-}nM^{-})$ | $k_{\text{un}}(s^{-})$ | $k_{\text{bleach}}(s^{-})$ | Number of states |
| --- | --- | --- | --- | --- |
| 4 | $1 \times 10^{-3}$ | $20 \times 10^{-3}$ | $10 \times 10^{-5}$ | 2 |
| 4 | $2 \times 10^{-3}$ | $20 \times 10^{-3}$ | $10 \times 10^{-5}$ | 2 |
| 4 | $5 \times 10^{-3}$ | $20 \times 10^{-3}$ | $10 \times 10^{-5}$ | 2 |
| 5 | $1 \times 10^{-3}$ | $10 \times 10^{-3}$ | $10 \times 10^{-5}$ | 2 |
| 5 | $1 \times 10^{-3}$ | $20 \times 10^{-3}$ | $10 \times 10^{-5}$ | 2 |
| 5 | $1 \times 10^{-3}$ | $50 \times 10^{-3}$ | $10 \times 10^{-5}$ | 2 |
| 6 | $1 \times 10^{-3}$ | $20 \times 10^{-3}$ | $10 \times 10^{-5}$ | 2 |
| 6 | $1 \times 10^{-3}$ | $20 \times 10^{-3}$ | $20 \times 10^{-5}$ | 2 |
| 6 | $1 \times 10^{-3}$ | $20 \times 10^{-3}$ | $50 \times 10^{-5}$ | 2 |
| 7 | $1 \times 10^{-3}$ | $20 \times 10^{-3}$ | $10 \times 10^{-5}$ | 2 |

**Table SI5:** Results from the Bayesian inference scheme. Summary of the results shown in all figures. The values are shown as *mean ± SD*.

| Figure | $k_{\text{bind}}(10^{-3} s^{-}nM^{-})$ | $k_{\text{un}}(10^{-3} s^{-})$ | $k_{\text{bleach}}(10^{-5} s^{-})$ |
| --- | --- | --- | --- |
| 4 | $1.031 \pm 0.012$ | $23.950 \pm 1.086$ | $8.800 \pm 1.418$ |
| 4 | $1.971 \pm 0.018$ | $23.500 \pm 0.680$ | $10.260 \pm 0.913$ |
| 4 | $4.588 \pm 0.030$ | $24.620 \pm 0.504$ | $8.446 \pm 0.674$ |
| 5 | $1.048 \pm 0.014$ | $9.623 \pm 0.612$ | $9.661 \pm 0.848$ |
| 5 | $1.031 \pm 0.012$ | $23.950 \pm 1.086$ | $8.800 \pm 1.418$ |
| 5 | $1.046 \pm 0.012$ | $45.650 \pm 1.586$ | $10.650 \pm 1.890$ |
| 6 | $1.031 \pm 0.012$ | $23.950 \pm 1.086$ | $8.800 \pm 1.418$ |
| 6 | $1.038 \pm 0.011$ | $22.470 \pm 1.455$ | $20.780 \pm 1.974$ |
| 6 | $1.014 \pm 0.012$ | $25.980 \pm 1.984$ | $43.150 \pm 2.852$ |
| 7 | $1.031 \pm 0.012$ | $23.950 \pm 1.086$ | $8.800 \pm 1.418$ |
| 7 | $1.008 \pm 0.021$ | $21.720 \pm 1.400$ | $11.940 \pm 1.937$ |
| 7 | $1.050 \pm 0.033$ | $23.480 \pm 2.720$ | $8.741 \pm 3.470$ |
| 8 | $2.774 \pm 0.077$ | $17.200 \pm 0.538$ | $0.725 \pm 0.355$ |
| 8 | $3.515 \pm 0.044$ | $14.460 \pm 0.234$ | $0.201 \pm 0.161$ |
| 8 | $8.110 \pm 0.067$ | $17.030 \pm 0.220$ | $0.419 \pm 0.207$ |

